# LC-Inspector: a simple open-source viewer for targeted hyphenated mass spectrometry analysis

**DOI:** 10.1101/2025.04.28.650946

**Authors:** Mateusz Fido, Etienne Hoesli, Elisa Cappio Barazzone, Renato Zenobi, Emma Slack

**Affiliations:** Department of Chemistry and Applied Biosciences, Swiss Federal Institute of Technology, Zürich, Switzerland; Department of Health Sciences and Technology, Swiss Federal Institute of Technology, Zürich, Switzerland; Department of Biology, Swiss Federal Institute of Technology, Zürich, Switzerland; Sir William Dunn School of Pathology, University of Oxford, UK; Basel Research Center for Child Health, Basel, Switzerland

## Abstract

The ubiquitousness of hyphenated mass spectrometry techniques across life sciences has made researchers around the world aware of their analytical power. However, the analysis of high-complexity mass spectrometry data remains virtually impossible for non-specialists. LC-Inspector is a standalone graphical user interface application for straightforward analysis of targeted mass spectrometry data, distributed under the MIT license and available free of charge for Windows and MacOS. The user can upload and process the data entirely locally on their machine by specifying the desired mass-to-charge ratios of the targeted compound ions and simply clicking “Process”. It allows for the processing of multiple files simultaneously, freely modifying and exporting graphs in real time, and calculating compound concentrations based on calibration standards. We show the versatility and applicability of LC-Inspector to different kinds of mass spectrometry data by analyzing a series of publicly available datasets from various samples recorded on instruments from multiple vendors and by different research teams.

## INTRODUCTION

Hyphenated mass spectrometry (MS) techniques have revolutionized the way analytical sciences shape today’s progress in science, industry and the clinic.^1–7^ Owing to the rapid development in instrumentation, spectrometers are now proficient at analyzing thousands of samples in record times and are capable of resolving many previously challenging isomeric and isobaric interferences. Liquid chromatography-mass spectrometry (LC-MS) has been at the forefront of analytical chemistry, becoming a leading technique and the analytical method of choice in drug discovery,^1,8,9^ pharmacology,^10^ the omics sciences^5,11,12^ clinical sciences,^7,13^ toxicology,^3,14,15^ and many others.^16–18^ Other branches of mass spectrometry have made strides in the fields of environmental sciences,^19^ structural biology,^20,21^ imaging,^22,23^ forensics,^24,25^ and virtually any other area where the sample’s structure, identity, composition or amount are of value.

Nevertheless, one of the consequences of the high throughput, precision and accuracy of high-resolution mass spectrometry measurements is that they tend to generate enormous datasets in the process. This “big data” problem requires mass spectrometrists or their collaborators to become skilled at conversion, processing, interpretation, and integration of this complex data or to employ additional bioinformaticians and data analysts for this purpose. The closed-source nature of the mass spectrometry vendor’s application programming interfaces (APIs) exacerbates this problem by making raw data inaccessible for processing without the usage of their proprietary tools. This leads to customer lock-in and promotes unhealthy dependency on the vendors’ products and workflow monopoly, a regrettably widespread practice.^26^

There have been several attempts at bridging the gap between generating highly complex mass spectrometry data and their processing and interpretation. Initiatives such as ProteoWizard^27^, XCMS^28^ or mzio’s MZmine^29^ have brought together large communities of applied MS users and are now widespread among research groups and companies alike. Since their inception, many related open-source chemometric tools,^30,31^ programming language libraries,^32,33^ frameworks and applications^34,35^ have been developed, further increasing the diversity of applied conversion, processing, and analysis methods.

Despite this, many challenges remain in the field of mass spectrometry data analysis. The multitude of instruments, possible data acquisition methods, workflows, and statistical methods used in life sciences make it unrealistic to apply a cut-and-dried analysis pipeline offered by most currently available software. Moreover, as MS data generally requires significant computational power, many analytical platforms require the user to upload their data onto an external server, further limiting their independence and shifting the trust to a third party, as the data is no longer processed locally on their workstation or personal computer.

LC-Inspector is a completely free-of-charge, fully local, and open-source program that provides a graphical user interface developed for viewing and analysis of mass spectrometry data. It integrates several Python libraries, enabling data visualization processing of different MS and LC data formats, and simple quantitative analysis. In this work, we present the main features of LC-Inspector and demonstrate its application to the preprocessing and analysis of several hyphenated mass spectrometry data sets.

## METHODS

### Application structure

The main graphical user interface is implemented via the model-view-controller architecture (MVC) using the PyQt6 framework and Python bindings for Qt (GPL v3 license).^36^ The main thread runs the user interface, while most of the resource-heavy computation is relegated to separate processes and threads using Python’s built-in multiprocessing, concurrency and futures modules. This minimizes UI-thread blocking, maintains user interactivity and makes maximum use of the available system resources. Files can be uploaded into LC-Inspector via the “Upload” tab of the main tab view Qt widget, with a custom list view supporting both drag-and-drop functionality and manual file browsing by displaying the OS-native file browsing dialog. The absolute paths are stored as a list of strings and converted into derivatives of the Measurement class upon starting the processing. The subclasses of Measurement define several methods for preprocessing the raw data, storing the processed data as properties, and enabling additional plotting modules.

### Supported input formats

The application accepts two types of data: text files (TXT and CSV file formats) containing chromatography data, and either mass spectrometry files (mzML) or peaklists with pre-assigned retention times, also in text format. mzML is an XML-based, vendor-agnostic file format that can be translated from any major manufacturer’s propriety mass spectrometry raw data file via tools such as MS Convert (ProteoWizard package).^27^ This allows for unified data sharing and analysis irrespective of the instrument and has been known for many years in the mass spectrometry community alongside other formats such as mzXML, mzMLb or HDF5.^37,38^

### Data processing

After loading the data, LC-Inspector performs the initial preprocessing. If operating in the LC/GC-MS mode, the chromatography and mass spectrometry data are processed concurrently. The chromatography data is parsed from any text file format containing columns of data convertible to Python floating-point numbers. Subsequently, baseline interpolation and correction are performed (described in the Results section). The program simultaneously reads the mzML files into memory, discarding any scans other than MS^1^. It then looks for the closest m/z values to the user-specified ion lists within a specific mass-to-charge range (3 ppm by default). It then rebuilds the extracted ion chromatograms for those m/z ranges, optionally connecting them to their chromatography retention times and automatically annotating the run with the desired MS-identified compounds.

### Plotting

The plotting takes place in a separate module and is based on pyqtgraph, a Python scientific graphics and GUI Library for Python.^39^ Pyqtgraph, as opposed to more popular and established libraries like matplotlib, maintains high user interactivity, enables real-time dynamic data processing such as log transformation, averaging, and fast Fourier transformation, and allows for exporting graphs in various formats. While not as mature as matplotlib (as of writing, 0.14.0 is the latest developer version of pyqtgraph), the library is comparatively lightweight, capable of performant plotting of millions of data points at the same time, dynamic panel docking and undocking, and embedding user-interactive and three-dimensional scientific graphs. It also provides limited capability for data export directly from the graph itself.

### Data export and further analysis

The export from LC-Inspector is a comma-separated values (CSV) file containing column-grouped data consisting of the filename, the traced ion’s name, its m/z value and an optional description, the ion’s retention time, integrated MS and LC intensities, and optional concentration. The format facilitates further processing or graphing via table-oriented software (e.g., R, pandas, GraphPad Prism, OriginLab Origin, Microsoft Excel) by saving data in vertical (long) format. The output of every analysis was subsequently processed using R.

## RESULTS AND DISCUSSION

### Working principle and design

The easy-to-use interface of LC-Inspector provides very simple access to the processing of raw MS data. The graphical layout of the application consists of a tab view, which is split into three sections: the Upload tab, where the user enters their data and provides initial parameters for the analysis. The Results tab, where they can view, modify and export the graphs plotted for each provided file, and the Quantitation tab is used to create calibration curves and calculate concentrations based on relative ion counts (Fig 1).

**Fig. 1.**
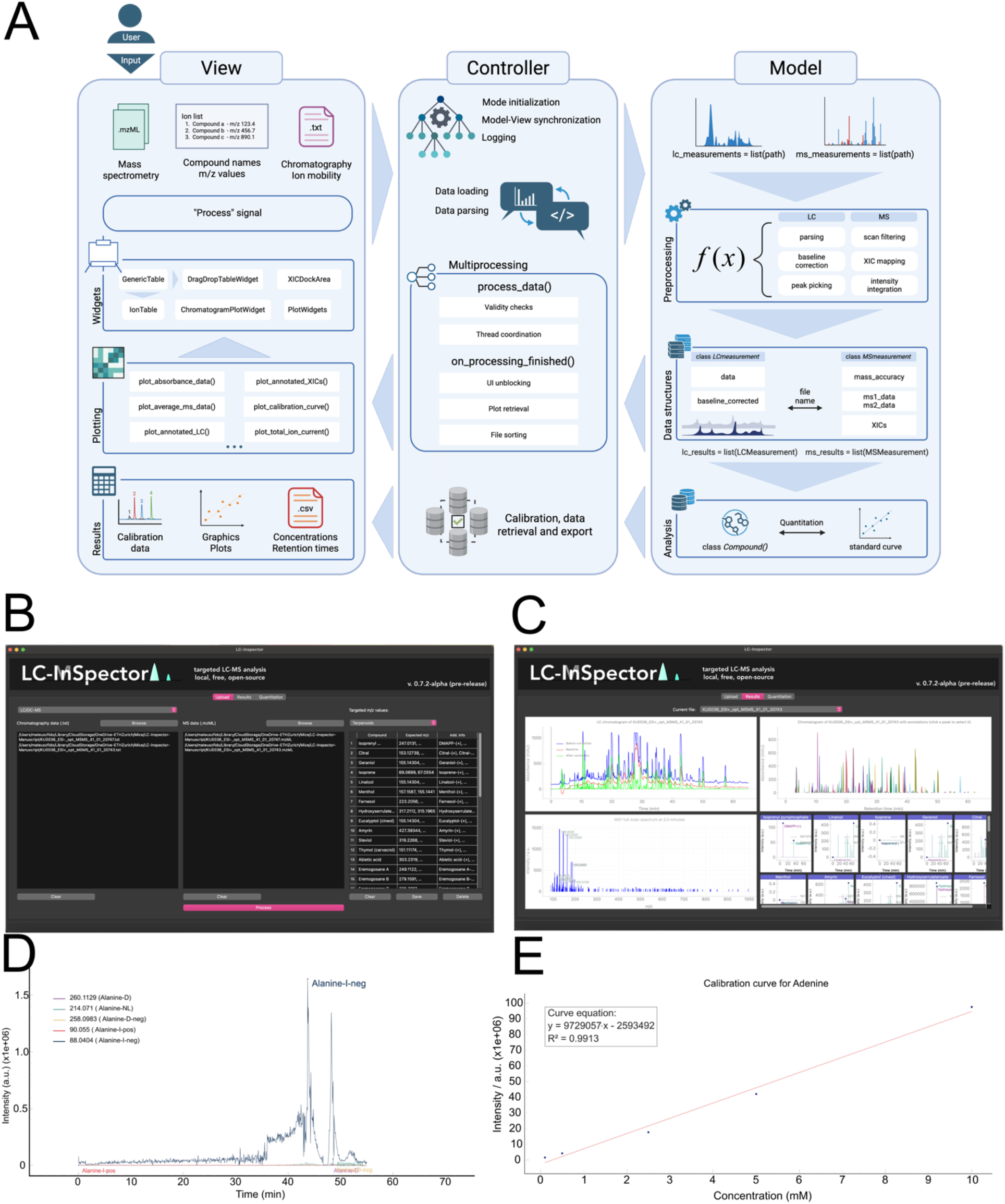
The graphical user interface of LC-Inspector. **A)** The application’s architecture and data handling model.^40^ **B)** The Upload tab in the LC/GC-MS mode, with the data table widgets and a sample targeted ion list chosen. **C)** The Results tab after processing of user-input data has finished. The four-panel layout in LC/GC-MS mode: the baseline-corrected LC/GC chromatogram (top-left), the chromatogram annotated with XIC-traced ions declared during the processing step (top-right), the mass spectrum respective to the current time on the chromatogram, changing dynamically when the user selects the desired time to view on the chromatogram in the top-left panel (bottom-left) and the extracted ion chromatograms of all the ions declared in the processing step (bottom-right). The bottom-right panels are expandable upon double-clicking them. **D)** An example of the expanded XIC panel containing plots of XICs for all the ions found for the given compound. **E)** An example calibration curve is obtained in the Quantitation tab.

Upon launching the application, the only available tab is the Upload tab, where the user can input their data. The main window area holds tables responsible for storing the absolute paths to MS data saved on the user’s filesystem, including shared and network disks. Each table widget supports drag-and-drop functionality for easy access, on top of being able to browse the operating system’s file manager with the “Browse” button.

LC-Inspector can work in three separate modes: “LC/GC-MS”, “MS Only”, and “Chromatography Only”. The user can choose the desired mode by using the combo box in the top left-hand corner of the Upload tab. The application then switches the input type, depending on the mode. The LC/GC-MS mode requires data in text file formats for the chromatography data, and the open mzML file format for the MS data. It then uses those data to display the chromatogram, the corresponding mass spectra at all retention times within the chromatogram, the chromatogram annotated with the respective *m/z* values and extracted ion chromatograms on which the annotation is based. The MS Only mode only requires mzML files for processing, treating the total ion current as a chromatogram-like representation of the time-resolved signal. The Chromatography Only mode requires the input of chromatography data, as well as retention time-based manual peak labels to display annotated data.

### Baseline correction

The core features of LC-Inspector, aside from the graphical interface, lie in its preprocessing techniques. One of the main examples is its chromatography baseline correction, which utilizes the Statistical Non-linear Iterative Peak clipping algorithm,^41^ similarly to the hplc-py package,^42^ where we saw good results for even highly variable LC run backgrounds, including negative absorbance values. Briefly, the algorithm works through approximating the baseline by applying the compressing LLS operator:

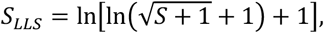

Where *S* is the original chromatography signal.

The algorithm then iteratively filters the compressed signal by selecting the minimum from the intensity at current scan time *t* and the mean of the signal at variable scan widths:

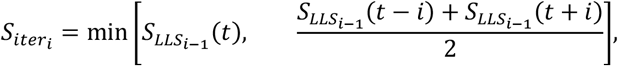

Where *i* is the current iteration. LC-Inspector uses a fixed default number of 20 iterations for all chromatography data, as we noticed this to be sufficient for most applications while keeping the computational preprocessing load relatively low. Finally, the signal is transformed back by transforming *S*_iter_ with the inverse of the LLS operator and subtracting from the original signal (Fig 2).

**Fig. 2.**
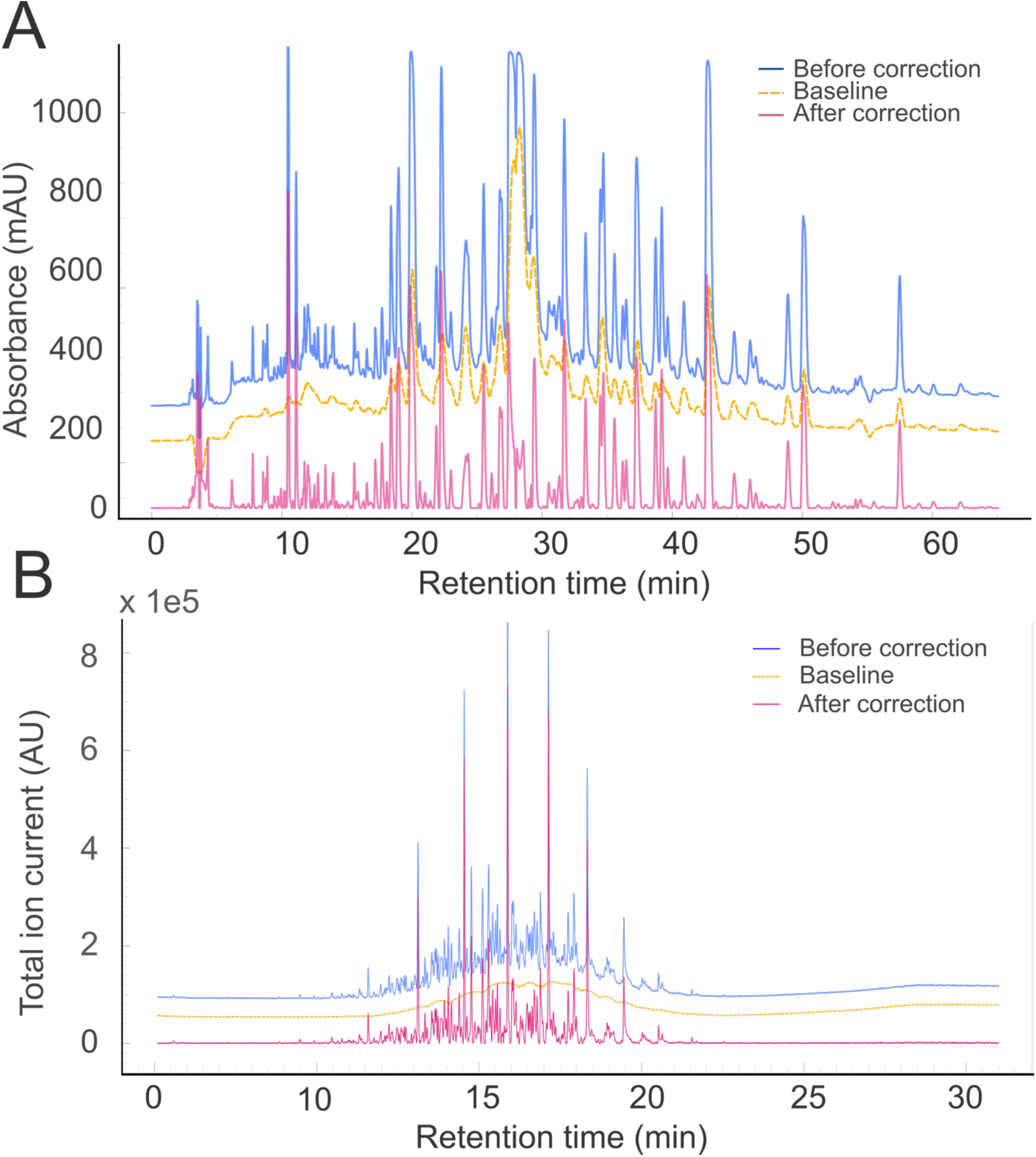
**A)** An LC chromatogram and **B)** a GC chromatogram before (blue, top) and after (red, bottom) applying the SNIP algorithm. The interpolated baseline is highlighted in yellow dashed lines (in between).

Figure 2A demonstrates the application of the SNIP algorithm on a very noisy spectrum obtained from the GNPS-MassIVE repository (MSV000088442, Agilent 1260 HPLC and Bruker micrOTOF-Q MS).^43^ The interpolated background is visibly contributing to the signal as the run progresses. Upon subtraction, the resulting corrected UV spectrum contains much sharper peaks and an effectively null baseline. Figure 2B presents the performance of the algorithm on a GC spectrum recorded with an Agilent 8890/5977B GC-MS system (NIST Public Data Repository, *<u>doi:10.18434/mds2-2601</u>*).^44^ The plots are directly exported from LC-Inspector (and recolored), facilitated by the underlying graphing library, which also allows extensive editing, removing, and transforming the plotted data in real-time.

### Quantification

Targeted mass spectrometry workflows often involve some kind of quantification, particularly within the omics sciences.^45–47^ To support this, LC-Inspector enables the user to select a subset of their input data as calibration data with known concentrations. Only one concentration value per file is supported, i.e., a single file represents a single calibrant data point, although an unlimited number of compounds is supported. Since this approach presupposes a large number of files, concentrations are imputed from the filename, though they can also be entered and modified manually. The user can choose to use either chromatography data or mass spectrometry ion counts for the calibration procedure, depending on the mode. Upon selecting the calibration files and clicking the “Calculate” button, a list of calibration curves is presented for every compound that was initially declared for tracing.

### Non-standard mass spectrometry

Considering the continuous rapid evolution and development of mass spectrometry, new hyphenated techniques emerge in the field on a regular basis. As a means of standardizing the input, LC-Inspector accepts mzML as its file format of choice, as it is ubiquitous in the field, and its conversion is supported by all the major vendors.^38^ This means that any mass spectrometry data that can be written in mzML can be analyzed, including flow injection and direct infusion techniques.

One example of such a method is secondary electrospray ionization mass spectrometry (SESI-MS), wherein a gaseous analyte collides with a charged electrospray, producing secondary ions.^48^ The method has gained popularity in the field of breath analysis, and general analysis of trace vapors. Figure 3 shows the application of LC-Inspector to SESI-MS data (ThermoFisher Scientific Q Exactive Plus) by using the MS Only mode. As the technique does not feature any chromatographic separation, the chromatogram is replaced by the total ion current, which serves as a global overview of the evolution of the signal across the time dimension.

**Fig. 3.**
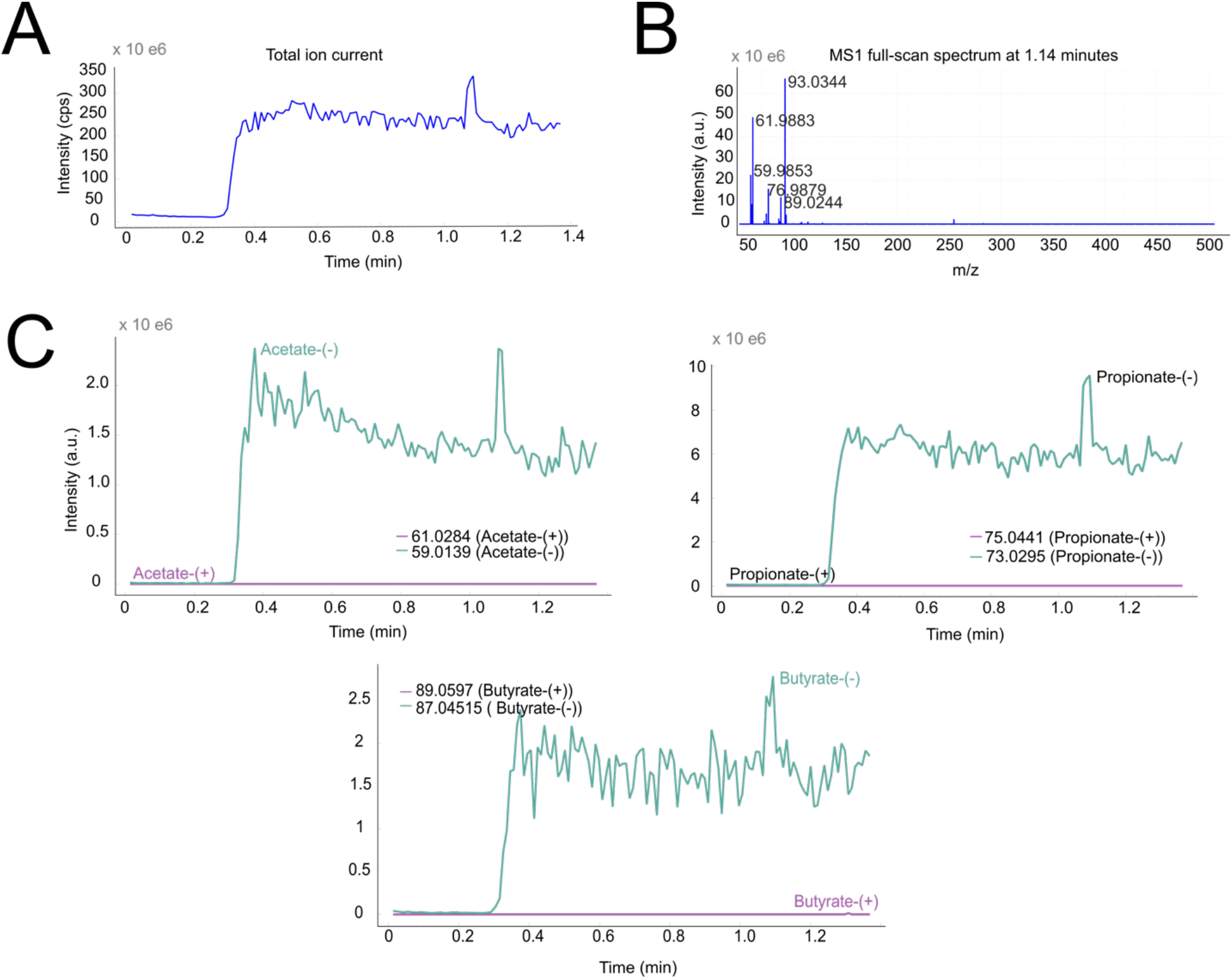
SESI-HRMS samples of human breath gas, which were stored in an ethylene vinyl alcohol copolymer sampling bag produced in-house, direct graphical output of LC-Inspector. **(A)** Interactive plot of the total ion current, which displays the underlying mass spectrum **(B)** when the user clicks on a corresponding scan time of the TIC. **(C)** Extracted ion chromatograms of short-chain fatty acids present in the exhaled breath samples (acetic acid, propionic acid, butyric acid) can be seen correlating with the total ion current.

As of today, there is no agreed-upon general analysis pipeline for the preprocessing and presentation of SESI-MS data. Most researchers default to analyzing and presenting data in the way of untargeted metabolomics, rarely investigating individual putative metabolites. LC-Inspector was created with targeted approaches in mind, facilitating the exploration of small molecules underlying the examined experimental conditions, preferably confirmed with further measurements (e.g., with tandem mass spectrometry, derivatization, standard spiking, etc.). Our hope is that researchers expand into targeted workflows and pursue confirmation of the traced metabolites, which our software aims to assist and expedite.

## CONCLUSION

We presented the free, local and open-source graphical interface software LC-Inspector, enabling quick and simple viewing and investigation of mass spectrometry data and discussed it in the scope of currently available technology. As MS becomes more widespread and common in laboratories throughout the world, parsing and evaluating the information it produces need to keep up with the technique’s prominence. In our view, this means removing barriers of entry wherever possible and making the analysis streamlined and available also for non-experts, which is why we hope this software helps adopt targeted workflows with minimal experience in mass spectrometry and data analysis. This could have broad appeal in the related fields, such as proteomics, lipidomics, glycomics, and metabolomics.

## SUPPORTING INFORMATION

LC-Inspector is distributed under the MIT license. The source code and executables for Windows and MacOS, as well as installation instructions, can be found free of charge on GitHub (https://github.com/MateuszFido/LC-Inspector). Version 0.3 (designed to be run on a high-performance computing cluster) can be downloaded from Zenodo (DOI: https://doi.org/10.5281/zenodo.13990448).

## ACKNOWLEDGMENTS

The authors would like to thank the early-stage users of LC-Inspector for helpful discussions and remarks when using the first versions of the software.

